## Supplementary Figures for "TranSPHIRE: Automated and feedback-optimized on-the-fly processing for cryo-EM"

### Supplementary materials

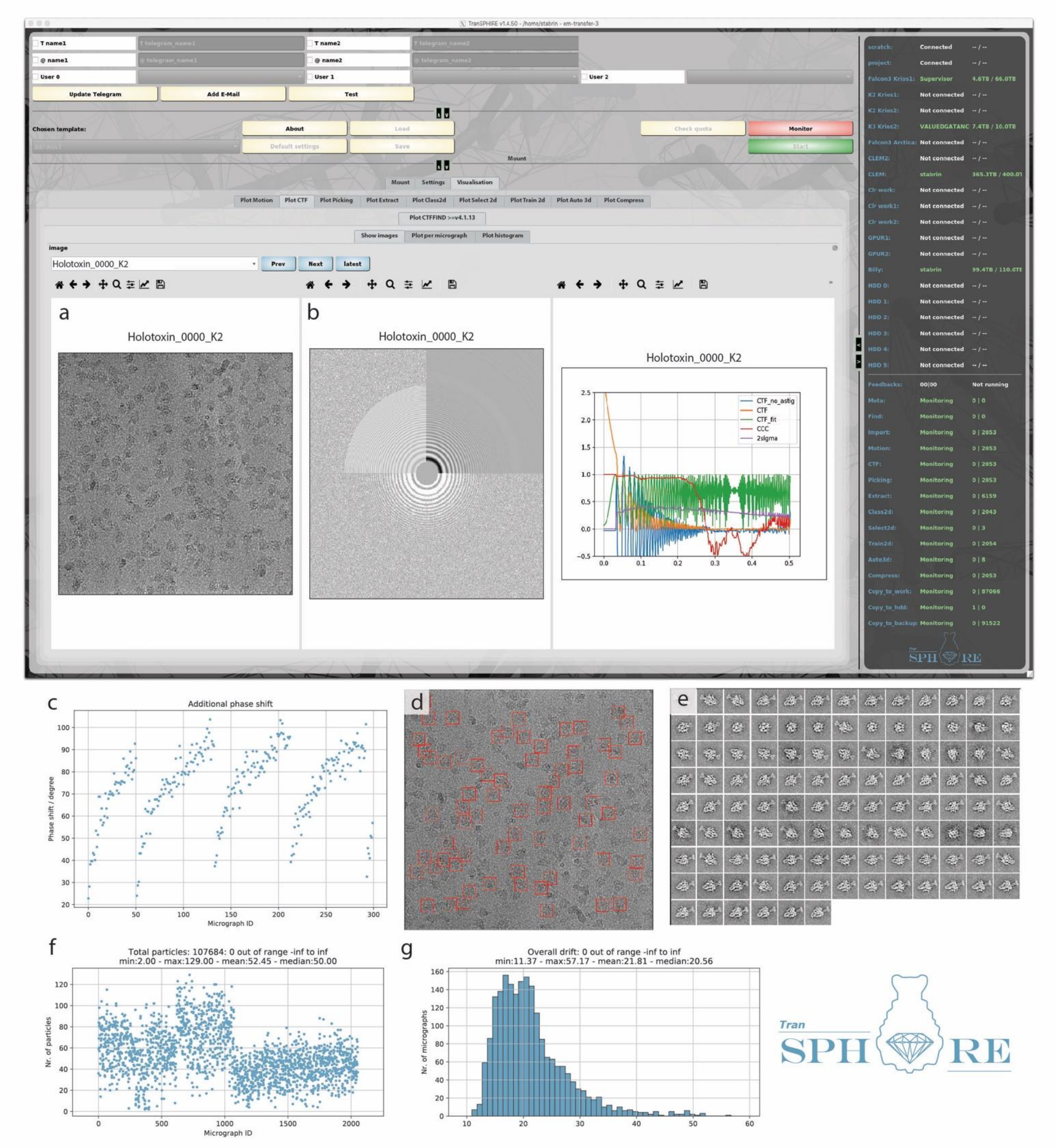


**Supplementary Figure 1. The TranSPHIRE graphical user interface (GUI).** The TranSPHIRE GUI is organized in tabs for initial setup, data processing settings, and live visualization of data acquisition and processing results. Visualization options include incoming micrographs (a), CTF fitting results (b), and phase shift (c), picking results (d), and 2D class averages (e). The shown phase shift development follows a logarithmic curve and helps experimentalists to decide when to switch to a new phase plate position (c). The shown picking result depicts the use of an optimized picking model, trained during runtime to only pick pore state particles (d). Output values can be plotted as either a scatter plots or histograms, as shown here for the total number of particles (f), and the overall drift per micrograph (g), respectively. This live monitoring enables an early evaluation of data quality during data acquisition and provides initial information about the protein structure.

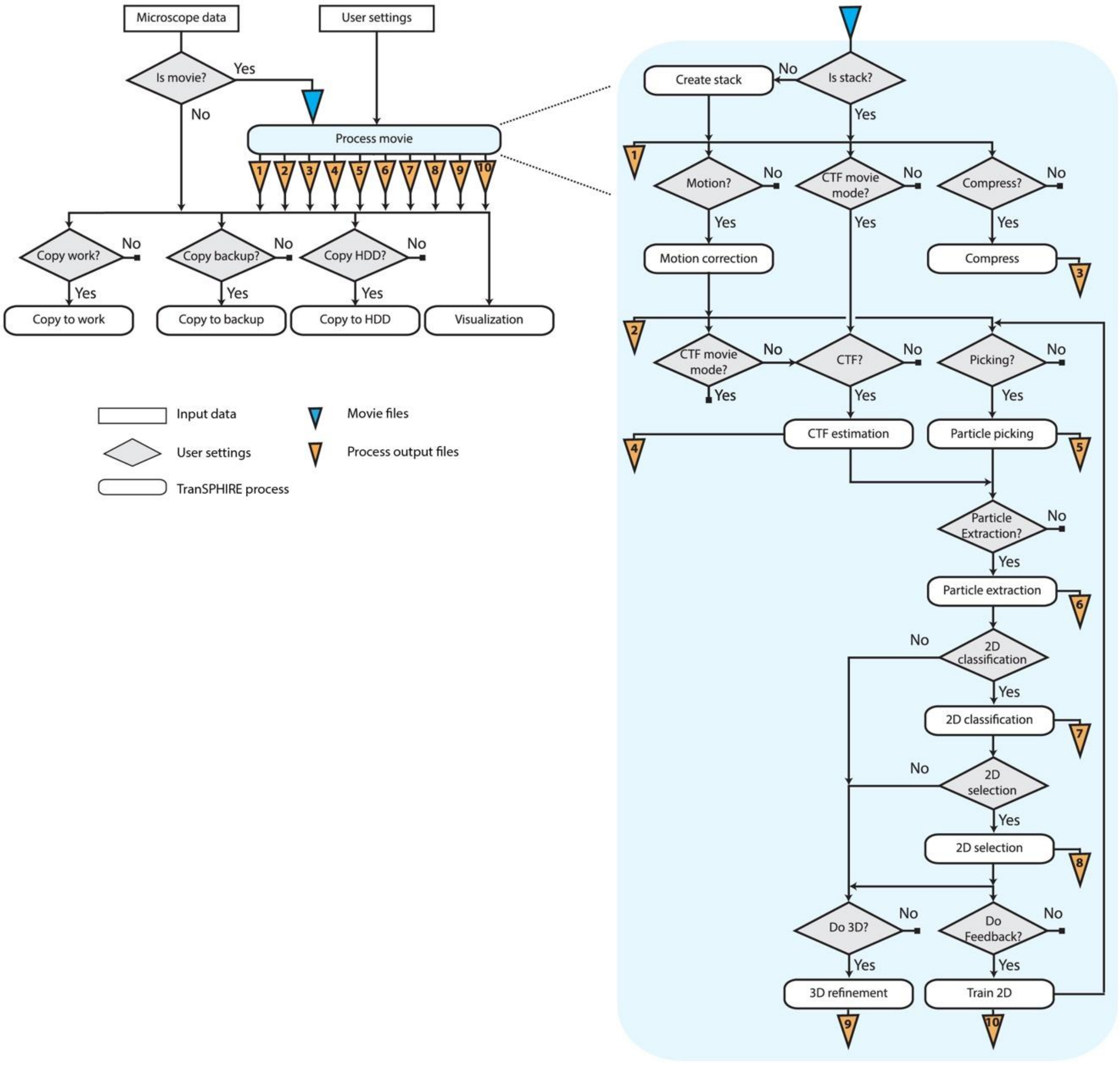


**Supplementary Figure 2. TranSPHIRE workflow.** Flow chart of the TranSPHIRE pipeline depicting sequentially executed processes below each other and parallel running processes next to each other. The workflow is highly adaptable allowing, for example, the binning of super resolution data during motion correction and CTF estimation. All inputs from the microscope, outputs from the involved processes, and additional statistics produced by TranSPHIRE are monitored and presented live in the

GUI. If specified, the processes “Copy to work”, “Copy to backup” and “Copy to HDD” create a copy of the results of each individual step to a workstation/cluster, a backup server, or an external hard drive, respectively.

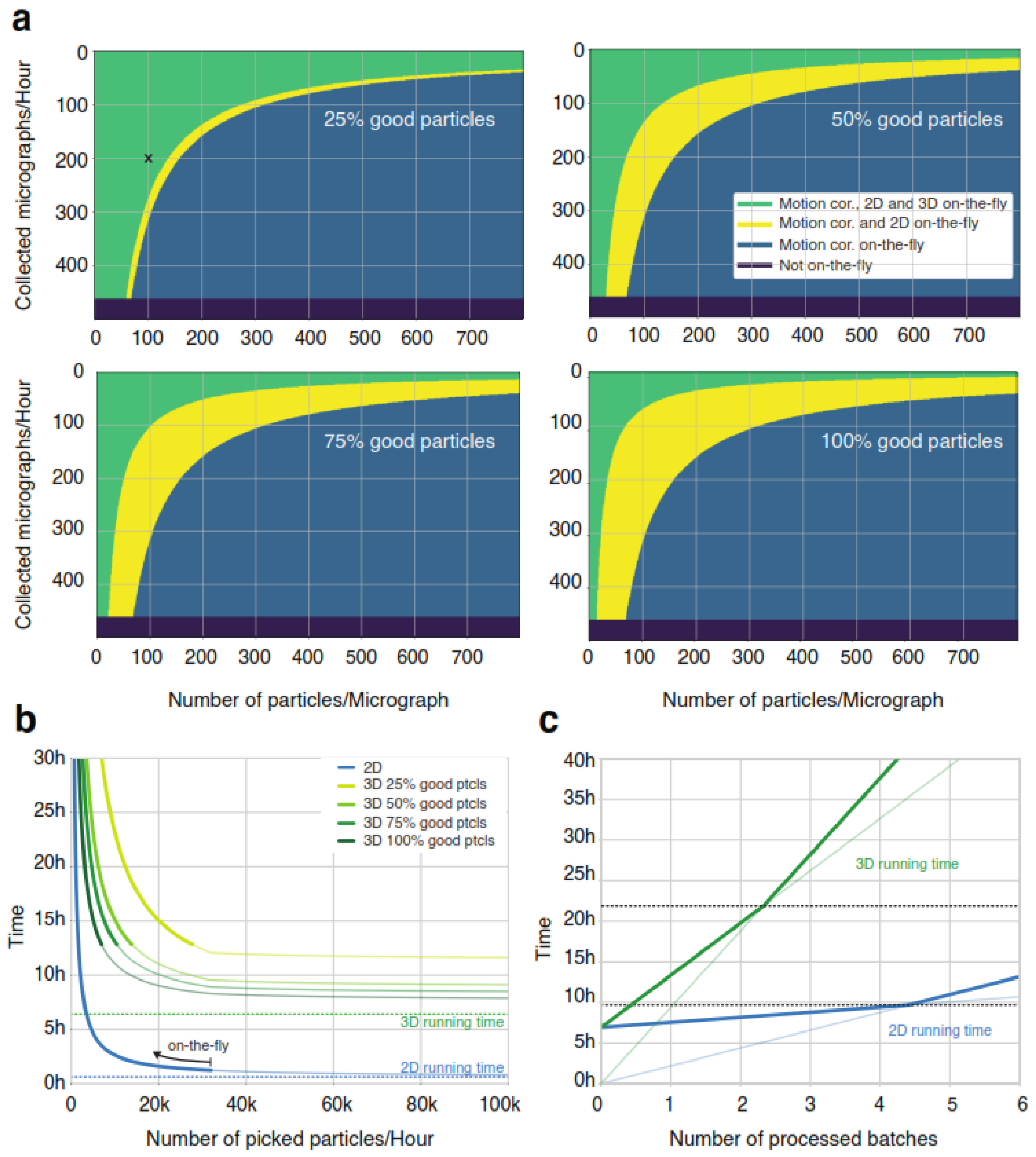


**Supplementary Figure 3. Benchmarks and timings for on-the-fly processing. (a)** Plot illustrating the dependence of on-the-fly processing on the average number of particles per micrograph (x-axis), the number of micrographs collected per hour (y-axis) and the percentage of “good” particles (displayed for 25%, 50%, 75% and 100%). TranSPHIRE can keep up with most routinely used data acquisition settings (green). In case of data collection with e.g. AFIS or a very dense particle distribution the hardware used in this paper might only allow on-the-fly processing up to 2D results (yellow) or motion correction (blue). Presented data are extrapolated from timings of the TRPC4 data processed in this manuscript. **Example:** *Supposing 25% of particles end up in classes labeled "good" by Cinderella (top left graph), the microscope provides 200 micrographs per hour, and 100 particles are picked on average per micrograph, then TranSPHIRE will provide motion correction, 2D processing, and 3D processing at on-the-fly speed* **(black cross)**. **(b)** Time needed to compute 2D class averages (blue) or a 3D reconstruction (shades of green) depending on the number of particles picked per hour and the percentage of "good" particles. Bold lines indicate the conditions where processing runs on-the-fly. Dashed lines indicate running times of 2D classification and 3D refinement (excluding the time needed to accumulate 20,000 particles for the 2D classification step and 40,000 useable particles for the 3D refinement step). **(c)** Number of batches needed to catch up with data acquisition after running the feedback loop. Data are simulated assuming an average of 115 picked particles per micrograph (50% of which labeled "good"), and an acquisition speed of 81 micrographs per hour. Processing of the data only starts once the feedback loop has finished (after ~7 hours) and TranSPHIRE requires time to catch up in order to process data on-the-fly. Dashed lines (black) indicate time points where processing has caught up for 2D classification (green), 3D refinement (blue), and starts running on-the-fly. Transparent lines indicate run times excluding the feedback loop. Timings are based on default settings (particle thresholds of 20,000 (2D) and 40,000 (3D) per batch).

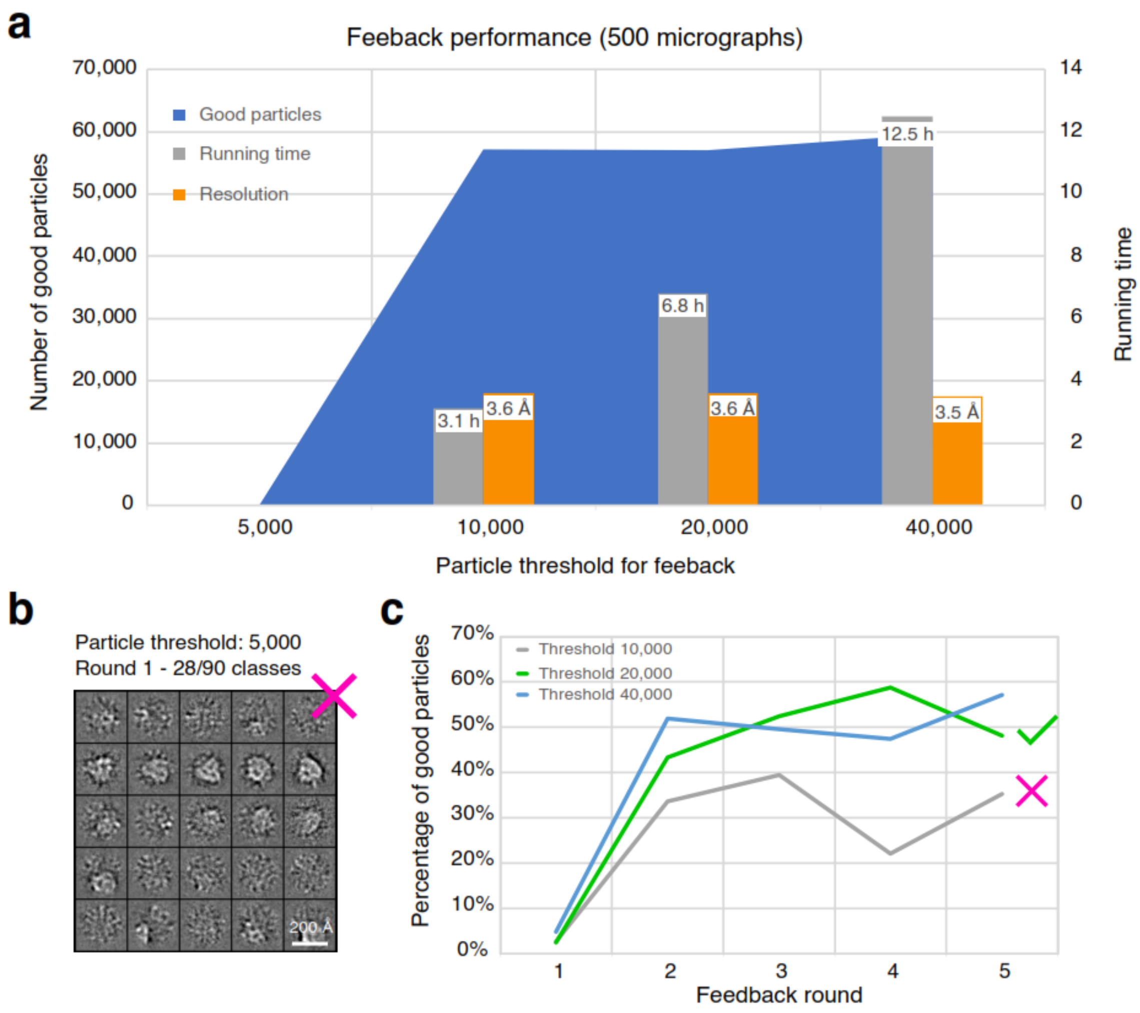


**Figure S4. Choosing an optimal particle threshold for the feedback.** To simulate a challenging data set, 90% of the picked particle coordinates were replaced by random coordinates on the micrograph prior to 2D classification. **(a)** Feedback performance measured as the total number of "good" particles (blue); the total running time of the feedback loop (grey); and the resolution of the final 3D reconstruction (green) in dependence of the chosen particle threshold. Data points for a particle threshold of 5,000 are missing, because the number of particles was insufficient for 2D clustering in round 1 (also see **b**). While both the resolution and number of good particles varies little, the running time increases considerably with an increases particle threshold. Thus, one has to balance the speed and stability when choosing a particle threshold for the feedback. (**b**) Representative 2D classes using a particle threshold of 5,000 illustrating that the number of particles is insufficient to generate high resolution classes**. (c)** Evaluation of the picking performance in dependence of the particle threshold. With a particle threshold of 10,000, the percentage of "good" particles fluctuates stronger and converges to a lower value compared to higher particle thresholds. As the performance is similar for particles thresholds of 20,000 and 40,000, a default value of 20,000 was chosen due to its lower run time.
